# A Scalable Framework for Closed-Loop Neuromodulation with Deep Learning

**DOI:** 10.1101/2023.01.18.524615

**Authors:** Nigel Gebodh, Vladimir Miskovic, Sarah Laszlo, Abhishek Datta, Marom Bikson

## Abstract

Closed-loop neuromodulation measures dynamic neural or physiological activity to optimize interventions for clinical and nonclinical behavioral, cognitive, wellness, attentional, or general task performance enhancement. Conventional closed-loop stimulation approaches can contain biased biomarker detection (decoders and error-based triggering) and stimulation-type application. We present and verify a novel deep learning framework for designing and deploying flexible, data-driven, automated closed-loop neuromodulation that is scalable using diverse datasets, agnostic to stimulation technology (supporting multi-modal stimulation: tACS, tDCS, tFUS, TMS), and without the need for personalized ground-truth performance data. Our approach is based on identified periods of responsiveness – detected states that result in a change in performance when stimulation is applied compared to no stimulation. To demonstrate our framework, we acquire, analyze, and apply a data-driven approach to our open sourced GX dataset, which includes concurrent physiological (ECG, EOG) and neuronal (EEG) measures, paired with continuous vigilance/attention-fatigue tracking, and High-Definition transcranial electrical stimulation (HD-tES). Our framework’s decision process for intervention application identified 88.26% of trials as correct applications, showed potential improvement with varying stimulation types, or missed opportunities to stimulate, whereas 11.25% of trials were predicted to stimulate at inopportune times. With emerging datasets and stimulation technologies, our unifying and integrative framework; leveraging deep learning (Convolutional Neural Networks - CNNs); demonstrates the adaptability and feasibility of automated multimodal neuromodulation for both clinical and nonclinical applications.

## Background

Closed-loop neuromodulation encompasses types of brain machine interfaces (BMIs) or human machine interfaces (HMIs) that monitor dynamic neuronal and physiological signals to optimize the timing and dosage of brain stimulation, as well as to tailor the stimulation parameters to a particular individual^1-3^. Closed-loop systems can be applied in clinical settings for several neurological disorders including age-related cognitive disorders or electrochemical disorders like drug resistant epilepsy. Nonclinical applications of such systems extend to health and wellness applications including attentional, behavioral, cognitive, or general performance enhancement. There are optimal and suboptimal times (“windows of opportunity”) for stimulation application, which derive from the dynamic nature of physiological/disease/performance states^4-11^. Conventional closed-loop approaches: 1) monitor a state (e.g., tremor, fatigue) or decode it from measured signals; 2) compare this current state to a desired target-state, producing an error signal which; 3) gates stimulation based on I/O models (of stimulation biophysics). Success of these approaches depends on the error signal reflecting optimal stimulation times and on stimulation-technology specific I/O models. Moreover, the tuning of such systems may integrate biases toward particular signal characteristics (e.g. triggered by signal amplitude at a given frequency) and selection of one stimulation type and may require ongoing feedback of outcomes which may not always be accessible.

Notwithstanding examples of such closed-loop approaches^12-20^, their generalization is limited by experiment-specific (subject-tuned) training, and specific assumptions on both meaningful error signals and stimulation mechanism of action. Many brain stimulation studies and clinical applications remain open-loop or minimally adaptive with single modality application, potentially applying stimulation interventions at suboptimal times relative to variations in targeted regions^21^; including most applications of transcranial electrical stimulation (tES). A preferred closed-loop algorithm - one that encourages deployment, development, and adoption – would minimize invasiveness and unnecessary stimulation; operate with any timescale suited to the targeted physiological/disease/performance state; and once programmed, would not require tuning with each participant’s ground-truth performance. Moreover, an extensively integrative, scalable, and generalizable system^22^ would employ data-driven optimization that can be incrementally trained from diverse data and any (multiple) stimulation modality (e.g., transcranial Direct Current Stimulation - tDCS, transcranial Alternating Current Stimulation - tACS, Transcranial Magnetic Stimulation - TMS, transcranial Focused Ultrasound Stimulation - tFUS) - amassing and integrating more datasets/modalities, as they become available, to enhance a convergent system’s capability. Such a system could be integrated into wearables for health and wellness technologies^23^ (attentional, cognitive, or performance enhancement etc.^24-26^) as well as expanded to clinical wearable neuromodulation technologies that facilitate and improve at-home treatment with primary neuromodulation interventions (i.e. chronic pain, fibromyalgia, multiple sclerosis, long COVID, drug resistant epilepsy etc.^27-33^) or used as a pharmacological treatment compliment (i.e. neurovascular modulation of the blood brain barrier^34^).

We present a flexible framework for designing and implementing closed-loop neuromodulation systems that utilize both deep learning techniques^35-37^ and workflows that avoid explicit state-decoding of stimulation modality (I/O model), and that gate stimulation based on a principle of responsiveness. Responsiveness depends on identifying epochs where a given stimulation modality will improve defined behavioral or physiological performance outcomes compared to not stimulating. These comparisons during training can be done with open-loop data while adapting ideas from causal inference such as potential outcomes comparisons^38-40^. To demonstrate and verify our framework we acquire and apply our analysis to the open-sourced GX dataset with convolutional neural networks (CNNs^35,41,42^) under data-driven optimization^43^. Leveraging electroencephalographic (EEG) and electrocardiographic (ECG) inputs, models identify periods of responsiveness, either positive or negative and map it to specific intervention types, which are then directly compared. We generate a framework-based decision process and demonstrate the identification of responsiveness with recommendations on intervention application (whether to apply stimulation or no stimulation) with given stimulation modalities (which specific stimulation type to apply). We explain how, provided datasets and a stimulation approach satisfying certain elements, our integrative approach allows for scalability and tunability across varied clinical and nonclinical neuromodulation applications.

## Results

### Scalable Closed-loop Neuromodulation Framework Architecture

The novel closed-loop framework is crafted from several aspects that make up the input (X(t)) to stimulation decision (S(t)) pipeline (**Figure 1c**). Input data (X(t)) is passed to a *Model Stack*, composed of an ensemble of independent models (numbered 1-M), each trained on a unique stimulation condition (in this case *No Stimulation, Stimulation Type 1*, and *Stimulation Type 2;* **Figure 1c**). Each ensemble predicts a *response* (i.e., class labels of an increase or decrease in behavioral or physiological outcomes) per its stimulation condition. These model outputs and confidences are compared under user-defined rules to produce stimulation decisions (*Stimulation Decision S(t)*). Importantly, the decision is based on responsiveness, namely an improvement in predicted performance (the potential outcome) with stimulation compared to predicted performance with no stimulation. A final stimulation decision (S(t)) is made where the stimulation type is applied, and the loop continues.

**Figure 1.**
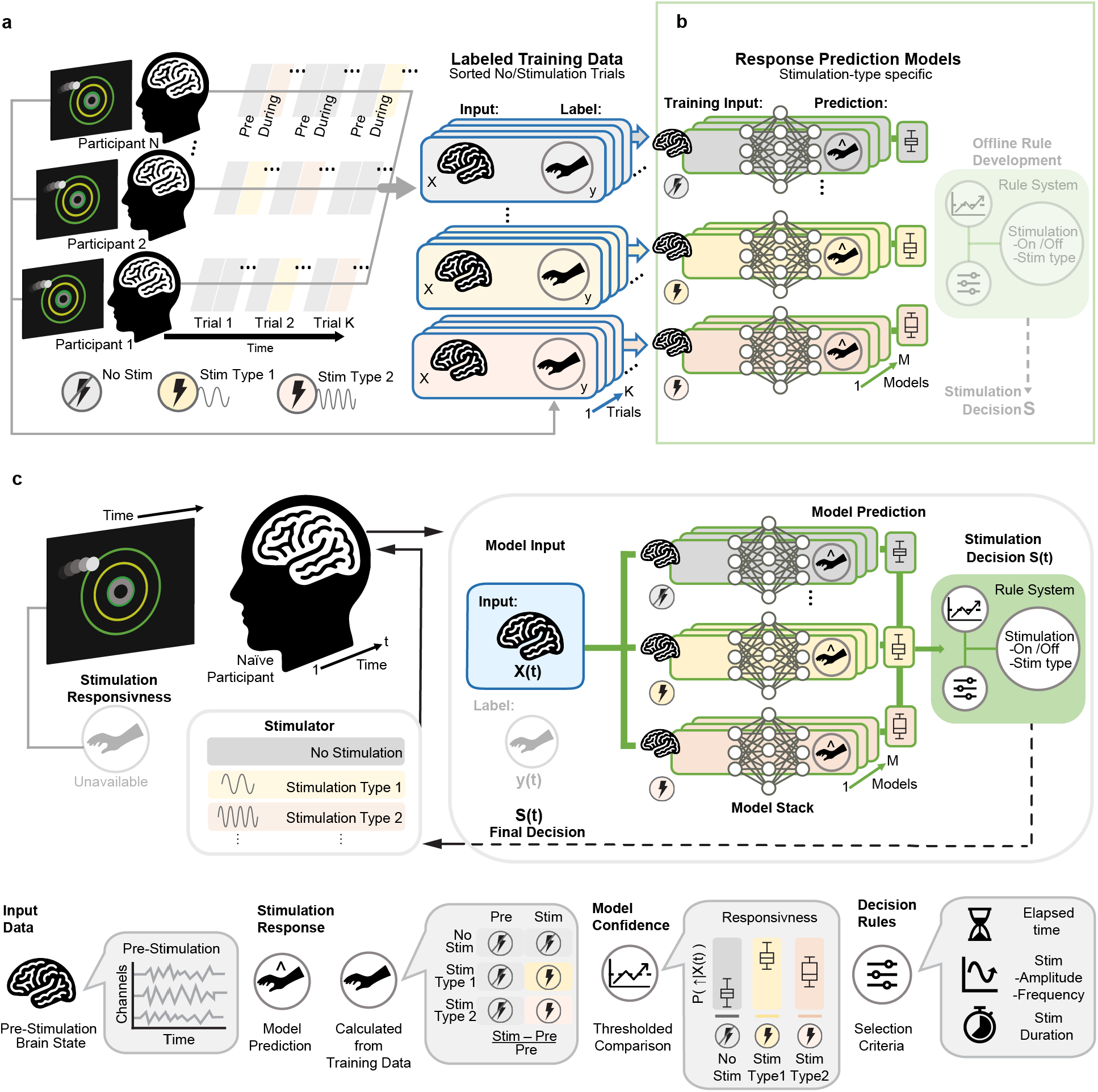
Novel framework for scalable closed-loop neuromodulation with deep learning. (**a**) Open-loop training data collection consists of multiple trials (*Trial*) across a cohort (*Participants*) where single trials record performance changes by stimulation (yellow and pink periods) as compared with no stimulation (gray). Note each stimulation intervention (e.g., Type 1: yellow, Type 2: pink periods) is preceded with a period of no stimulation (gray) where biomarkers (brain state) are collected. Training data can be arranged into biomarkers pre intervention (training *Input X*) and stimulation response (*Labels* y). (**b**) Multiple varying model architectures (spanning model space *M*) ingest multi-channel and multi-sample input data. Note, each set of model ensembles are trained on different intervention types, which allows for interchanging intervention types as needed. Next, the independent selection of stimulation decision (S) rules allows for the incorporation of extraneous factors such as stimulation cost. (**c**) In a closed-loop deployment, biomarker measurements (X(t)) from a novel participant are passed to each ensemble (e.g. No Stimulation, Stimulation Type 1, Stimulation Type 2) of the *Model Stack*. Each ensemble yields a performance prediction with *Model Confidences* for the condition it was trained for, which are then compared together to support specific stimulation *Decisions Rules*. The stimulation decision (S(t)) selects among the stimulation candidates or no stimulation, is delivered to the participant, and the loop continues. Note, in deployment, ground truth (y(t)) data are not required, and models can be independently updated as more training data becomes available. The implementation of a stackable ensembled workflow lends advantages of scalability, agnosticism to (multiple) stimulation modalities (e.g., TMS, tFUS), without explicit biomarker decoding or I/O models, or need for personalized ground-truth performance. See bottom key for definition of symbols.

At the point of training, application of our framework for data-driven closed-loop neuromodulation using deep learning starts with designing an appropriate open-loop stimulation experiment (**Figure 1a**) and selecting appropriate deep learning model architectures (**Figure 1b**). The outcomes of these stages serve to verify the feasibility of the given implementation (described below) and ultimately (with additional data collection / training) drive a closed-loop implementation (**Figure 1c**).

The behavioral task used in training sessions determines the performance target for the closed-loop system. The stimulation modalities used in training are the options available to the closed-loop intervention, based on biomarkers used within the training data. A simple way to design open-loop trials is with a continuous performance metric, measuring acute changes in performance in response to stimulation, and with biomarkers collected prior to the stimulation (and in the absence of stimulation) in order to compose a causal structure for the stimulation intervention to predict responsiveness.

Our framework does not require that a biomarker predict (decode) performance before or in response to stimulation. Further, whereas conventional closed-loop systems compared decoded brain state/performance against a desired level in order to trigger stimulation; here we require only that the biomarker-derived features indicate a brain state where simulation is likely to improve outcome. For example, if a closed-loop algorithm that inadvertently resorts to detecting sleep in EEG and uses that as an indicator of task performance, then this is far from ideal; since in this case the algorithm would chose to stimulate only when sleep is detected, continue to do so if the participant stays asleep, and not necessarily stimulate when stimulation would benefit the participant to change the outcome measure. In contrast, stimulation responsiveness identifies brain states suggesting both poor performance without stimulation and improved performance with stimulation, similar to potential outcomes frameworks.

At the point of implementation (**Figure 1c**) the input data, denoted as X(t), is provided to the models, and the input data labels (y(t)) are not needed since models are already trained. The input data can consist of a segment of time series data (such as EEG or ECG), sampled at a particular time (t), that should be a biomarker of responsive brain state (**Figure 1c**). There are several model ensembles which form a *Model Stack*. Each model ensemble is previously trained to precise responses to its designated stimulation type (*Stimulation Type 1, Stimulation Type 2…*) and one ensemble trained to identify response to no stimulation (**Figure 1c**, *No Stimulation*). Within each model ensemble set, each model (models 1 - M) can have different architectures or compositions to extract different characteristics of input data, however, all models within an ensemble set should be trained on the same stimulation modality and training data. For example, the *No stimulation* model ensemble set can contain multiple convolutional neural network (CNN) models that each have different kernel sizes or different layering architectures.

The main advantage of model stacking is that it allows for interchangeability and flexibility between ensemble sets within the *Model Stack* meaning interventions can be added or removed as needed without the need for robust retraining. For example, the *Stimulation Type 1* can be completely removed from the system leaving behind only the *No Stimulation* and *Stimulation Type 2* stacks. Similarly, a new model ensemble can be added to the *Model Stack* as needed, to include a new stimulation type. This not only reduces training time but allows for utilizing different cohorts for training data (e.g. varied experiment durations, varied number and rates of stimulation, and both open-loop or closed-loop sessions), provided that their input data type and performance metrics match the *Model Stack*.

A further aspect of our pipeline involves comparing the outputs from each ensemble of the *Model Stack* to make a stimulation decision (S(t)). Importantly, this decision is based on the principle of responsiveness which compares the performance predicted with each stimulation type and no stimulation. For example, if the *No Stimulation* model ensemble predicts that a participant’s response will not increase if no stimulation is applied, and the *Stimulation Type 1* ensemble set indicates that the participant’s response will improve with this type of stimulation, then this indicates responsiveness to *Stimulation Type 1* and suggests it should be applied. On the other hand, if the *No Stimulation* model ensemble predicts that a participant’s response will increase with no stimulation applied, and the *Stimulation Type 1* ensemble predicts a smaller improvement, then this indicates no responsiveness to *Stimulation Type 1* and suggests no stimulation should be applied.

Next, the determination of which stimulation type (S(t)) to apply based on *Model Stack* outputs, is application specific, and defined by decision rules, which can compare all potential outcomes. One method would be to consider an average or majority vote among each model ensemble set (where predictions can be binary outcomes – improvement or no improvement) as well as some metric of confidence with each prediction (e.g., comparing the prediction probability of improvement across all stimulation types, where prediction probabilities are averaged within each ensemble set). The decision rule, as an independent stage, is application specific. For example, if the *No Stimulation* model ensemble predicts that a participant’s response will moderately increase if no stimulation is applied, and the *Stimulation Type 1* ensemble set indicates that the participant’s response will significantly improve with this type of stimulation, a decision rule may still limit stimulation based on factors like the cost of stimulation (i.e., tolerability, power consumption, elapsed time since last stimulation bout etc.).

In training a closed-loop algorithm (from open-loop data) the notion of responsiveness needs to be stipulated: we are concerned with identifying biomarker-based features that predict when stimulation is likely to produce an improvement in performance compared to no stimulation. A condition where performance would increase or decrease regardless of if stimulation is applied, can be considered neutral from the perspective of the stimulation value. A condition where performance would increase with stimulation and decrease without stimulation application (i.e., due to vigilance decrements, inattentiveness, or sleepiness), is considered positive from the perspective of the stimulation value. A condition where performance would decrease with stimulation and increase without stimulation application, is considered negative from the perspective of the stimulation value. Responsiveness-based gating supports the preferred system feature of minimized unnecessary stimulation (“light touch”).

There is no stipulation that datasets have a homogenous format (e.g., duration of experiment or number of test stimuli) supporting scalability, and any distinct stimulation modality can be developed in parallel supporting integration. Predicates for our framework is the existence of (and identifying) at least one fixed stimulation dose that is broadly effective, and the identification of generalized brain/physiological states that track the time of sensitivity to the ascribed dose - together representing responsiveness. We speculate such features are more likely to exist for interventions with limited focality and where targeted performance is dynamic (on the timescale of mins to sec). Use of non-invasive approaches is consistent with our suggestion of a preferred platform for scalable training and deployment - such as tES / EEG considered in the next section.

### Framework Application to Exemplary Dataset

Our framework for closed-loop neuromodulation can be applied to a wide range of applications with use-case specific implementation. We illustrate one application which is based on acquisition and analysis of the open-sourced GX dataset, containing continuous EEG, ECG, EOG and behavioral metrics (**Figure 2b**) in response to High-Definition tES with varied doses (**Figure 2c-d**)^44^.

**Figure 2.**
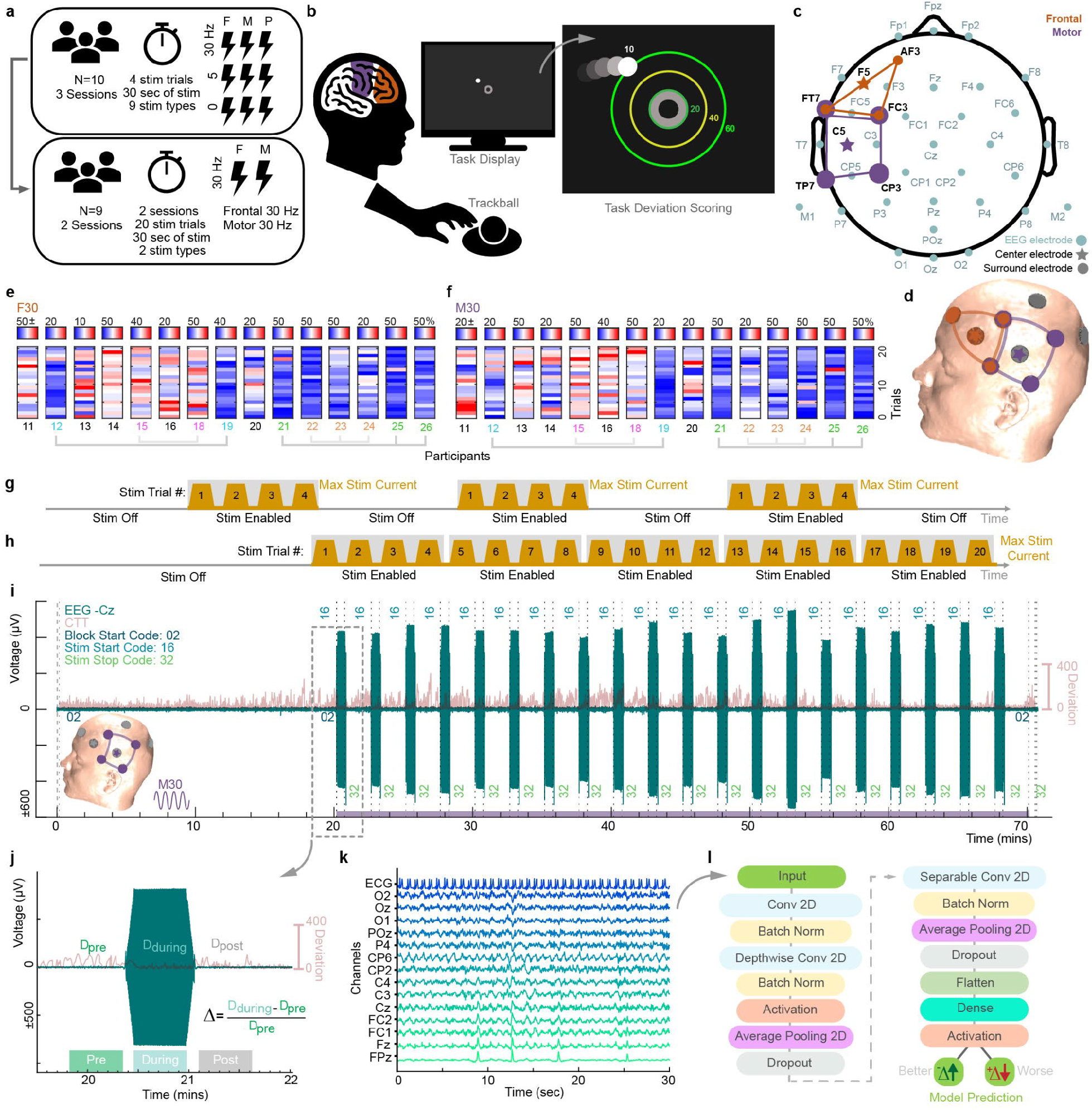
Dataset summary and model architecture. (**a**) The GX dataset was collected in two phases or experiments. Experiment 1 consisted of a parameter space exploration where 9 different stimulation conditions were tested. This was followed with Experiment 2 which tested two parameters (F30 and M30) from Experiment 1 on a different participant cohort with 20 trials of each parameter. (**b**) The behavioral task for the GX dataset consisted of a 70-min compensatory tracking task (CTT) where participants used a trackball to keep a moving circle at the center of the screen. The radial distance or deviation from the center of the screen at each frame determined participant’s behavioral performance. (**c**) EEG (light blue) and stimulation sites for frontal (orange), and motor (purple) stimulation. (**d**) MRI-derived head model used to visualize stimulation electrode placement. Trial-wise percent improvement derived from change in deviation for all participants in Experiment 2 for (**e**) F30 and (**f**) M30. Trial-wise implementation of stimulation trials for (**g**) Experiment 1 repeated three times for each participant to cover all 9 parameters and (**h**) Experiment 2 repeated two times for each participant to cover 2 selected parameters (F30 and M30). (**i**) EEG and CTT timeseries data for one session for participant 24 where M30 was applied. Insets indicate stimulation location. (**j**) Trials were segmented from each session into 30 sec pre, during, and post stimulation. The percent change in CTT deviation (Δ) was calculated from averaged CTT data in the course of the pre and during stimulation periods. See panel **e** for trial-wise and participant-wise results. (**k**) A typical segment of input training data consisted of 15 channels (14 EEG and 1 ECG) X 3000 samples (30 secs). (**l**) A CNN architecture was utilized to as a classifier to predict response indicated by changes in the percent change in deviation (Δ) during the CTT (*Better* or *Worse*).

For this implementation biomarkers of response consisted of 32-channel pre-stimulation EEG and ECG, and labels of response included a behavioral compensatory tracking task (CTT) where participants’ goal was to use a trackball to keep a dynamic cursor-controlled circle at the center of the screen at all times. Lower radial ball deviation from the center of the screen indicated good task performance (**Figure 2j**). Stimulation conditions included 9 different combinations of stimulation location (Frontal, Motor, Parietal) and frequency (0, 5, 30 Hz), denoted with the first letter of the location and the frequency number (i.e., Frontal stimulation at 30 Hz as F30; **Figure 2a**). In acquiring the GX dataset, Experiment 1 was used as a parameter space exploration to identify stimulation conditions that produced the largest degree of behavioral improvement and demonstrate an open-loop effect (**Figure 2a**). Experiment 1 served an important function in framework implementation (**Figure 1a**) since a detectable open-loop effect is a prerequisite. Frontal (F30) and Motor (M30) stimulation at 2 mA and 30 Hz, were selected from Experiment 1 and reimplemented in Experiment 2 with more trials and a different cohort (**Figure 2g-h**).

Once an open-loop stimulation effect on performance is established, models are trained to predict when stimulation would most effectively change performance. In our case, we utilized the data acquired during Experiment 2 as our input training data since we could effectively extract response from individual trials where segments of no stimulation (or pre stimulation) preceded segments of stimulation. We utilized the percent change in average task cursor deviation (Δ) or distance from the center of the screen at each screen frame (∼16 Hz), from pre to during stimulation (**Figure 2b and j**) as our response label (see **Figure 2e** for trial-wise and participant-wise results) and the pre-stimulation EEG and ECG (15 channels X 3000 samples) as our predictive brain states to train each of the individual models (**Figure 2k**). For our no stimulation comparisons, data where no stimulation was applied was divided similarly to that of the stimulation portions of data (see **Figure 2i** from 0-20 mins). Each pre-stimulation trial was labeled as *Better* or *Worse*, where class labels were binned identifiers calculated from the percent change in behavioral performance (CTT deviation; **Figure 2j** Δ) during stimulation compared to the pre-stimulation period. A negative percent change in deviation (-Δ, compared to pre stimulation) was classed as *Better*, whereas a positive change in deviation (+Δ, compared to pre stimulation) was classed as *Worse*. In terms of model architecture, a modified CNN (EEGNet architecture^42^) was used with differing input kernel lengths (**Figure 2l**).

### Framework Verification with Exemplary Dataset

The acquired GX dataset was used to present and verify a proof-of-concept of the application of our framework^44^. Data from Experiment 2 met our framework criteria (for training data) of an open-loop effect of stimulation and CNN models were selected to be applied to the dataset. For our framework verification and proof-of-concept, data from Experiment 2 were selected for further analysis in order to maintain homogeneity with experimental intervention timing and maximize the number of training trials.

For each stimulation condition (No Stimulation, F30, M30) two models, each with differing kernel lengths were tasked with producing binary classifications of trials of pre-stimulation input data (14 EEG and 1 ECG channels X 3000 samples). Each of the 6 models produced prediction accuracies of trial-wise binarized change in deviation (*Better*: a CTT deviation less than pre stimulation period, *Worse*: a CTT deviation more than pre stimulation period) above 50%, indicating that models were able to effectively ingest minimally processed input data (EEG and ECG) and that the input data across stimulation conditions contained predictive information on responsiveness (**Figure 3d-f**). Models 1 and 2, for the No Stimulation condition reached respective cohort testing (816 total trials; Better/Worse ratio: 1.0503) accuracy of 73.5% and 73.2%, and misclassifications of 26.5% and 26.8% (**Figure 3d**). Similarly, for the F30, both Models 1 and 2 reached cohort testing (120 total trials; Better/Worse ratio:2.6364) accuracy of 67.5%, and misclassifications of 32.5% (**Figure 3e**). For M30, Models 1 and 2 reached respective cohort testing (120 total trials; Better/Worse ratio: 3.2857) accuracy of 71.7%, and 69.2%, and misclassifications of 28.3% and 30.8% (**Figure 3f**).

**Figure 3.**
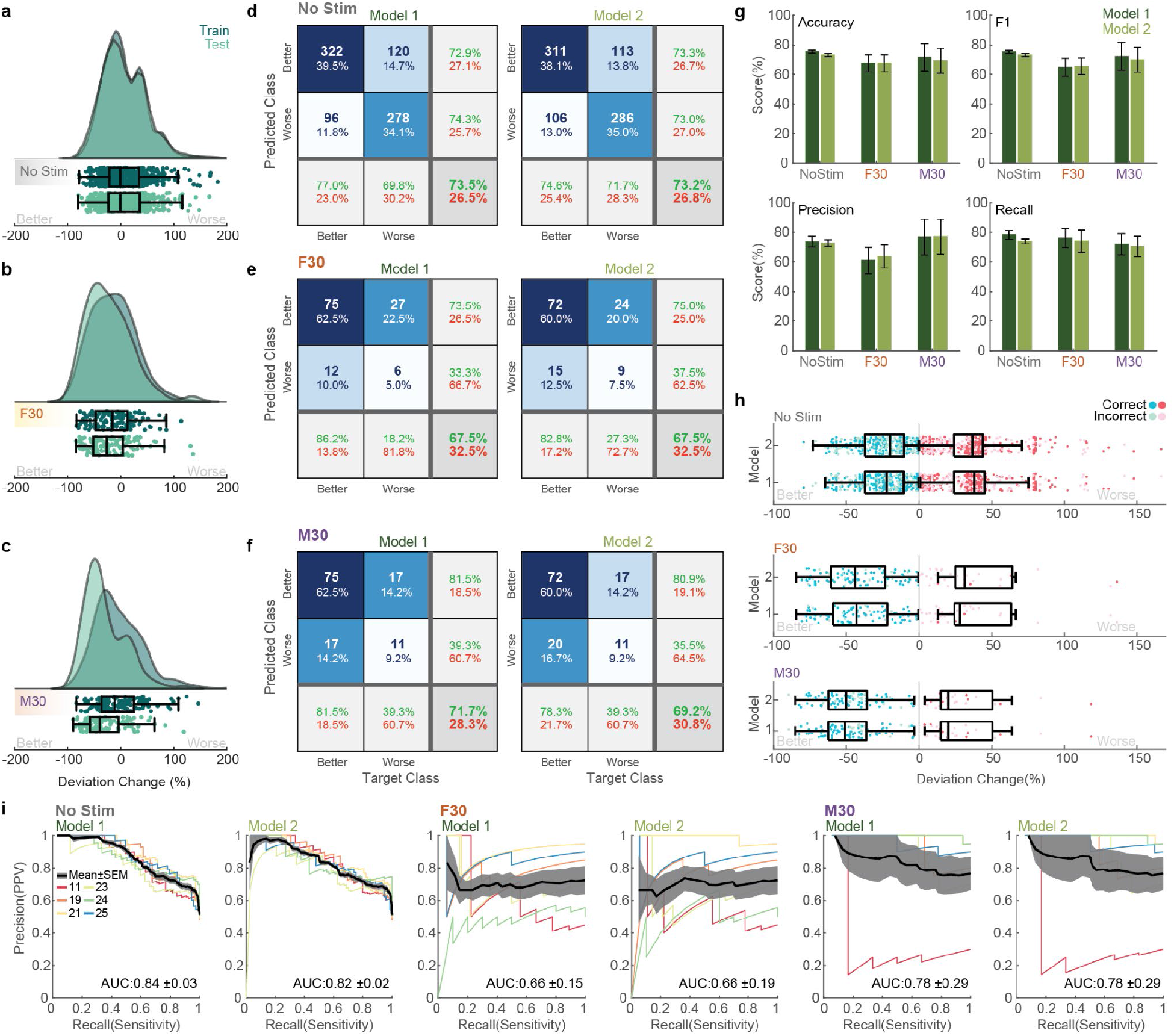
Data label distribution and model prediction metrics. Distribution of training and testing data used for (**a**) No Stimulation, (**b**) F30, and (**c**) M30 stimulation conditions. Confusion matrices for Model 1 and 2 for (**d**) No Stimulation, (**e**) F30, and (**f**) M30. (**g**) Scores for Models 1 and 2 in terms of accuracy; and weighted scores for precision, recall, and F1 with SEM. (**h**) Correct and incorrect classifications for Models 1 and 2 with their respective percent change in deviation. Boxplots are indicated for correct classifications only. (**i**) Precision-recall curves across test data input type and models with the area under the curve (AUC) computed for the average precision recall across participants with SEM.

Due to the open-loop effect of stimulation across stimulation conditions (**Figure 3a-c**), training and testing data suffered from class imbalances between both classes (*Better* and *Worse*). Weighted (by class frequency) metrics of precision, recall, and F1 were therefore calculated to better reflect model performances (**Figure 3g**). Aforementioned metrics along with the area under the precision-recall curves (PR-AUC; **Figure 3i**) were computed as the mean and standard error of the mean (SEM) across participants and trials (N=6, Stim: 20 and NoStim:136 trials for each participant) in the test set. All model metrics are summarized in **Table 1** and **Figure 3i**.

**Table 1:** Summary of model performance metrics (mean± SEM) for each of the two models for No Stim, F30, and M30.

|  | No Stim |  | F30 |  | M30 |  |
| --- | --- | --- | --- | --- | --- | --- |
|  | *Model 1* | *Model 2* | *Model 1* | *Model 2* | *Model 1* | *Model 2* |
| **Accuracy** | 75.4±1.0 | 73.2±1.0 | 67.5±5.7 | 67.5±5.7 | 71.7±9.4 | 69.2±8.5 |
| **F1** | 75.2±1.1 | 73.2±1.0 | 64.9±6.2 | 65.6±5.7 | 72.1±9.3 | 70.0±8.5 |
| **Precision** | 73.6±3.7 | 72.8±2. | 60.9±8.8 | 63.7±7.8 | 76.9±12.1 | 77.1±11.9 |
| **Recall** | 78.3±3.1 | 73.9±1.6 | 76.1±6.5 | 74.2±7.6 | 72.0±7.1 | 70.4±7.0 |
| **PR-AUC** | 0.84±0.03 | 0.82±0.02 | 0.66±0.15 | 0.66±0.19 | 0.78±0.29 | 0.78±0.29 |

Predictions for the No Stimulation condition had a relatively balanced correct classification between both classes and correct classifications were widely distributed over percent changes in deviation (**Figure 3h**). For F30 and M30 stimulation conditions, correct predictions were skewed to the major class (*Better*), where a higher percentage of the major class was correctly predicted as compared to the minor class (*Worse*).

These results indicate that models within ensemble sets (e.g., Model 1 and Model 2 would be an ensemble for *No Stimulation*) for all stimulation conditions utilized, can effectively identify out-of-training responsive trials. These predictions (for each stimulation type) can then be compared directly, post-prediction, to determine the appropriate stimulation decision and stimulation application of each trial (see **Figure 1c** and **Figure 4**).

**Figure 4.**
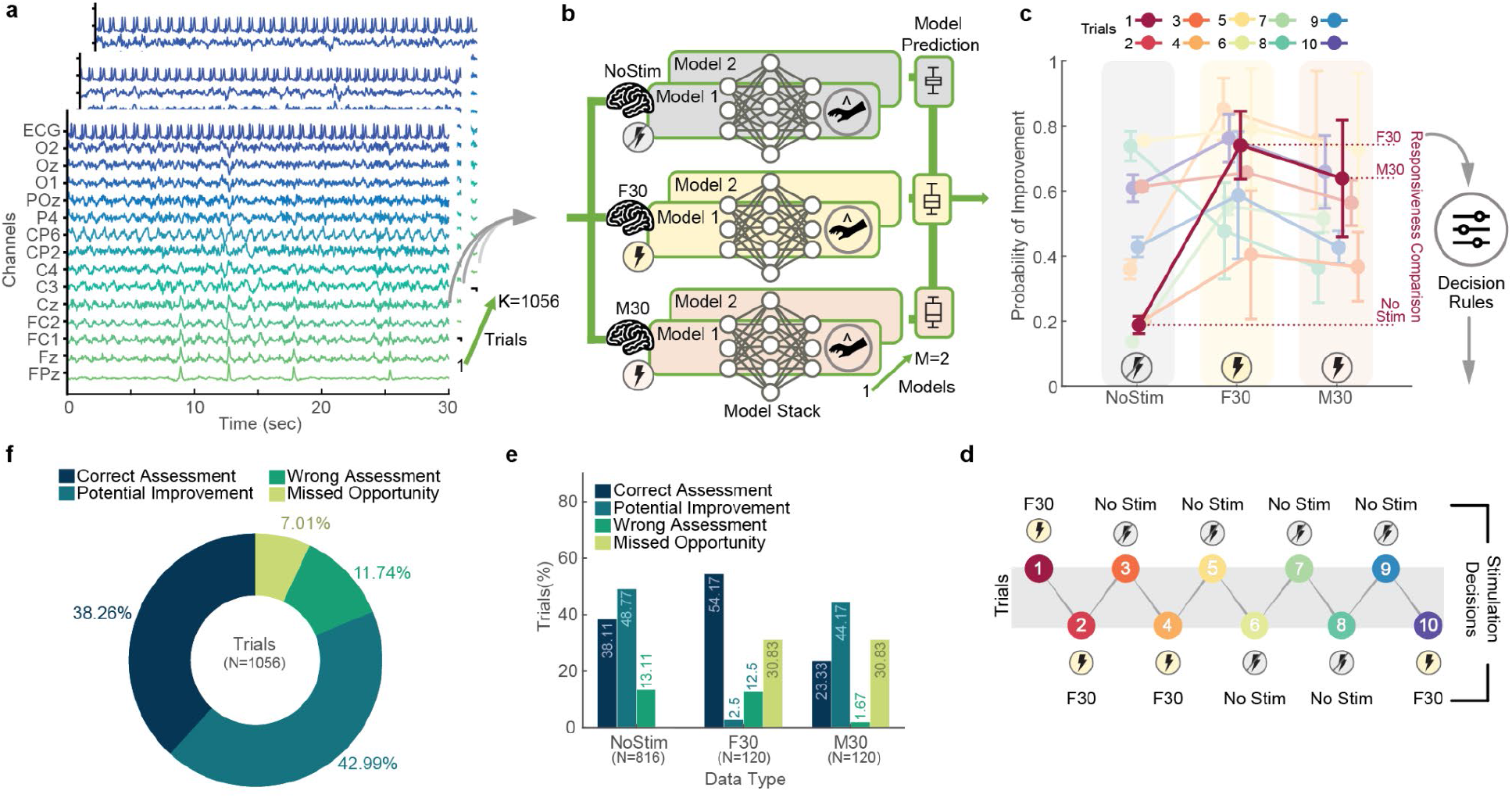
Simulation of closed-loop responsiveness predictions. (**a**) A total of 1056 input test data trials (15 channels X 3000 samples) were individually passed to all 6 models. (**b**) The predictions from all 6 models were aggregated and compared post prediction. (**c**) Model predictions (with SEM) for 10 exemplary trials where the probability of improvement (probability of trial classed as *Better*) is averaged over predictions for each respective model (i.e., predictions from Models 1 and 2 for NoStim, F30, and M30). Using simple decision rules, model predictions are compared. (**d**) Stimulation decisions for 10 example trials in **c**. (**e**) To assess simulation outcomes, stimulation decision predictions of test trials where no stimulation, F30, and M30 stimulation were applied were divided into four classes: correct assessment, potential improvement, wrong assessment, and missed opportunities. (**f**) These four classes were then aggregated across all test data.

Once trained all models can then be combined to produce responsiveness predictions. To demonstrate responsiveness predictions, comparisons, and stimulation decision making; all test trials were individually passed as inputs to an ensemble of all 6 models (**Figure 4a-c**). Model predictions were aggregated and averaged, and responsiveness comparisons (**Figure 4c**) were made followed by a simple decision rule, where the maximum probability of improvement (or potential outcome) across stimulation intervention types (above a threshold of 0.65) was selected as the stimulation decision. **Figure 4c-d** demonstrates the responsiveness comparisons and stimulation decisions for 10 exemplary trials. In terms of the responsiveness comparisons with our implementation, a decision rule algorithm first considers if the average NoStim probability of improvement is more than 0.65. If yes then the stimulation decision is NoStim, if not then F30 and M30 are compared directly to check that both their probabilities of improvement are more than 0.65 and that one average probability is more than the other, leading to either F30 or M30 being chosen as the stimulation decision. If these criteria are not met the stimulation decision reverts to NoStim, inline with our “light touch” approach to stimulation.

To assess the responsiveness predictions, stimulation decisions were classified into correct assessments, potential improvements, wrong assessments, or missed opportunities. This was done for each test set data type as well as all test set trials (**Figure 4e-f**). Trials classed as correct assessments were instances where the stimulation decision matched the factual label of the given stimulation intervention that was applied and showed an improvement in behavioral performance. Potential improvements indicated trials where the stimulation decision was an intervention type that could have resulted in a behavioral improvement based on open-loop effects. Wrong assessments were trials where the stimulation decision was an intervention that was known to produce a given decrement in behavioral performance, given factual trial labels. Missed opportunities were trials where the stimulation decision was to apply no stimulation, even though factual trial labels indicated that there would be behavioral improvements with a given intervention type (i.e., F30 or M30).

## Discussion

This work presents a novel framework for implementing data-driven, integrative, closed-loop stimulation that is scalable in terms of stimulation modalities and training datasets; agnostic to stimulation modalities and dataset format; and omits the need for ground truth performance data in its application stage. Leveraging the acquired and open-sourced GX dataset^44,45^ as base brain stimulation training data and deep learning techniques^35,41,42^, we demonstrate applicability and a proof-of-concept. We show that with this structured technique, minimally processed neural and physiological input data can be used to effectively identify conditions anticipating stimulation responsiveness. Using responsiveness identification, our framework’s flexible decision process for intervention application identified 88.26% of trials as correct applications, showed potential improvement with varying stimulation types, or missed opportunities to stimulate, whereas 11.25% of trials were predicted to stimulate at inopportune times. Overall, in a closed-loop system, these identified states of responsiveness, within our framework, combined with mutable decision rules (e.g., causal inference) predict when and which stimulation will optimize performance/outcomes.

Our framework leverages tools from prior closed-loop techniques but provides special benefits in overall implementation. Agnosticism to modality allows any stimulation modality (e.g., electrical^46,47^, light^48,49^, sound^3,50,51^) to be incorporated, moreover using open-loop training data. Although multimodal stimulation is rare, our framework accommodates the use of multi stimulation types and opens both clinical and research avenues of exploration for these types of stimulation applications^52^. As such we avoid an explicit I/O model (hypothesized mechanism). Our framework parallelizes stimulation predictions across modalities allowing integration into a single controller. The expandable training stage is distinct from closed-loop implementation, omitting the need for ground truth performance in the target participant. Together these support scalability. Our approach allows for selecting a timescale for updating predictions, that would be informed by the time-course of the stimulation and performance change.

Our framework revolves around the concept of responsiveness. This circumvents the reliance of closed-loop systems to explicitly decode a latent brain state (or performance) to compare a target condition, with the resulting error triggering stimulation (based on a separate I/O model). Rather, responsiveness predicts how a given stimulation modality will change performance, which can be compared to expected performance without stimulation (or with another stimulation modality). Application-specific decision rules can then be implemented, for example based on the confidence of prediction or the costs of stimulation. These decision rule classifications can be adjusted to accommodate stricter (as in our “light touch” to stimulation applied here) or more flexible classifications, leading to potential reductions in outcome variability; and can be contrasted with open-loop stimulation applications (stimulation applied without informed timing or responsiveness), which can have larger outcome variability and probabilities of stimulation application at inopportune times. Once fully defined, our framework can be implemented to make autonomous stimulation decisions upon deployment.

As demonstrated here our framework leverages deep learning techniques, namely CNNs, which have been shown to be particularly effective for ingesting and classifying timeseries signals like EEG and ECG; however, our framework is agnostic any one specific deep learning technique and can integrate several different types of models. Inspired by the cytoarchitecture and processing pathways within the visual cortex, CNNs utilize spatial and temporal computations on input data in order to create hierarchical representations. These hierarchical feature representations can be explicitly examined to produce interpretable learned spatial and temporal filters and have exhibited increased performance with smaller models as compared to larger ones^53^. Similar to other deep learning techniques applied to neural and physiological (EEG, ECG, electromyography - EMG, photoplethysmography - PPG etc.) data classification^35^, such as recurrent neural networks (RNNs)^54^, generative adversarial networks (GANs)^55^, and Long Short-Term Memory (LSTMs)^56-58^; CNNs too require large amounts of labeled training data, an issue typically addressed with data augmentation as applied here. Indeed, there currently exists a dearth of large open-sourced neuromodulation datasets, however with advances in data modeling and augmentation^59^ techniques (diffusion models, variational autoencoders – VAEs, GANs etc.^60-64^), transfer learning^65,66^, and improved data integration and sharing infrastructure with data harmonization^67^, these deficits will be addressed.

Our framework can be applied to any recording and stimulation approaches, invasive or noninvasive, while adopting a data-driven approach to circumvent biases related to brain state and performance or the mechanisms of action of stimulation. The application of our approach therefore spans existing neuromodulation technologies and emerging interventions such as 1) using wearables^23^ to guide invasive stimulation (e.g. spinal cord stimulation for pain^68^ or deep brain stimulation for essential tremor^69^); 2) invasive^70-74^ and non-invasive^20,75^ closed-loop neuromodulation to enhance memory and cognition 3) invasive^76-80^ and noninvasive^81-83^ closed-loop neuromodulation for movement disorders and pain; and 4) EEG^84,85^ or heart rate^86,87^ guided TMS. The implementation here with the GX dataset is exemplary as a demonstration of our proposed framework and validating our approach. With additional experimental datasets and techniques, the performance of our framework becomes expandable with potential increases in predictive performance and wider applicability.

## Methods

### Dataset Description

To explore the flexibility of our proposed framework we acquired and utilized the open-sourced GX dataset^44,45,88-91^. The dataset is one of the largest concurrent tES, EEG, ECG, and behavioral datasets, contains over 68 hours of EEG data recorded during a continuous tracking task developed by Makeig et al^92,93^. Data used from the GX dataset consisted of a total of 19 participants (7 females, 12 males). Participants ranged in age from 19-43 (mean age 29.10±6.75 years) and were recruited from the New York Metropolitan area. All experimental procedures were reviewed and approved by the Western Institutional Review Board and all procedures were conducted in accordance with the ethical guidelines set forth by the Declaration of Helsinki in 1964 and its later amendments. All participants were financially compensated for their participation. Data collection and procedures are detailed elsewhere^44^ and is briefly detailed below.

#### Experimental Design

The GX dataset was collected in two main phases or experiments. The first experiment was a parameter space mapping experiment that explored different combinations of applying tES at different scalp locations and different stimulation frequencies. In total, Experiment 1 explored 9 stimulation conditions, which consisted of 3 scalp locations: Frontal, Motor, and Parietal; and 3 stimulation frequencies: 0, 5, 30 Hz (**Figure 2a**). Each stimulation condition was applied across 4 trials in each participant, with each bout of stimulation lasting 30 secs with a 5 sec ramp up and 5 sec ramp down (**Figure 2g**). In total, data from 10 participants was collected for Experiment 1, where the 9 stimulation conditions were applied across three 70 min sessions (3 conditions per session). At the conclusion of Experiment 1, the stimulation combinations of Frontal 30 Hz (F30) and Motor 30 Hz (M30) were selected as the best candidates to examine in Experiment 2. These were selected based on their open-loop behavioral effect and similar scalp sensation.

For Experiment 2, F30 and M30 were applied to each participant over 20 trials, where each trial consisted of 30 sec of stimulation with a 5 sec ramp up and 5 sec ramp down (**Figure 2h**). The application of these stimulation conditions was broken up across two 70 min sessions, where one stimulation condition was applied per session. In total 9 participants completed Experiment 2 and 4 participants were asked to return either once or twice to repeat both experimental sessions.

#### Behavioral Task

The behavioral task consisted of a continuous compensatory tracking task (CTT)^92^. The goal of the task was to keep a moving circle at the center of the screen at all times (**Figure 2b**). The circle was endowed with oscillatory and dampening forces, allowing it to be responsive and in constant motion. Participants controlled the location of the circle with their dominant hand using a trackball. The radial distance from the center of the screen at each time point (∼16 ms) indicated vigilant performance over the task duration. The task was performed continuously over each of the 70 min sessions, in a dark room. Some participants were offered foam earplugs for ambient sound attenuation. During task performance, participants were left undisturbed even if they exhibited hypnagogic states or fell asleep.

#### EEG and Physiology

EEG data were acquired using a 32-channel wired EEG cap (ANT Neuro, Hengelo, The Netherlands) using the 10/10 international system (**Figure 2c**). Within the Waveguard cap, 29 plastic holders (Soterix Medical Inc., New York, USA) were interleaved to allow for the application of electrical stimulation. Electrolyte gel (SignaGel, Parker Laboratories Inc., New Jersey, USA) was used between the electrode scalp interface for both EEG acquisition and electrical stimulation and was placed using blunt tip syringes (15-gauge; Cortech Solutions Inc., North Carolina, USA). Signals were grounded at AFz, references to CPz and sampled at 2 kHz. The amplifier voltage range was set to 1 V peak to peak with a bandwidth of 0-520 Hz. EEG electrode impedances were monitored and adjusted to <20 kΩ prior to data acquisition. Lead I ECG and EOG were acquired from bipolar snap electrodes. For ECG electrodes were placed across the chest below participants’ left (anode) and right (cathode) clavicles, whereas for EOG electrodes were placed at the left (anode) and right (cathode) outer canthus of participants’ eyes.

#### HD-tES

During both Experiments 1 and 2 electrical stimulation was applied for 30 secs with a 5 sec ramp-up and ramp-down time. Electrical stimulation was applied through 9 Ag/AgCl sintered ring electrodes (Soterix Medical Inc., New York, USA) at standard EEG 10/10 locations using a current controlled current source (MxN 9-channel high-definition transcranial electrical stimulation stimulator; Soterix Medical Inc., New York, USA). Stimulation electrodes were placed in plastic holders (Soterix Medical Inc., New York, USA), where electrolyte gel (SignaGel, Parker Laboratories Inc., New Jersey, USA) was used to interface between electrodes and scalp^94^.

For both experiments, electrodes were placed in an HD-MxN ring configuration where one electrode was placed at the center and between 3-4 were used as outer electrodes. For Frontal stimulation the return electrode was placed at F5, whereas the surrounding electrodes were AF3, FT7, and FC3. For Motor stimulation the return electrode was placed at C5, whereas the surrounding electrodes were placed at FT7, FC3, CP3, and TP7. For Parietal stimulation the return electrode was placed at CP3 whereas the surrounding electrodes were placed at C5, C1, P1 and TP7. Stimulation was only applied at one location per trial.

Stimulation waveform frequencies consisted of 0, 5 and 30 Hz. Waveforms were applied as either a monophasic DC (0 Hz) or as biphasic sinusoidal waveforms (for 5 or 30 Hz). With DC stimulation the center electrode was used as the cathode whereas the surrounding electrodes were used at anodes.

### Data Preparation and Deep Learning Models

#### Dataset and Preparation

All data preprocessing was conducted with custom scripts in MATLAB (2019b, Mathworks, Natick, USA) and Python (3.9, Python Software Foundation). Associated toolboxes included the EEGLAB toolbox^95^, ROAST^96,97^, Raincloud plots^98^, Tensorflow with Keras (version 2.7.0)^99^, Pandas (version 1.3.4)^100^, Numpy (version 1.20.3)^101^, Seaborn (version 0.11.2)^102^, Scikit-learn (version 1.0.1)^103^, and SciPy (version 1.7.1)^104^.

EEG and physiological data were minimally preprocessed prior to utilization. Data were baseline corrected between a 0-25 sec period at the start of each recording (for each 70 min session). Each 70 min session was divided into 30 sec trails. For Experiment 1 and 2, during the stimulation enabled (Stim Enabled) periods data were divided into trials based on stimulation triggers (**Figure 2i-j**). All data were portioned into three 30 sec periods (**Figure 2j**): 30 secs before stimulation was applied (*Pre*-Stim.), 30 secs during stimulation (excluding ramp-up and ramp down intervals; *During* Stim.), and 30 secs after stimulation was applied (*Post* Stim.). Similarly, the stimulation off periods (*Stim Off*) were sequentially divided into trials with a 15 sec overlap between the Pre and Post Stim periods. EEG during stimulation was excluded from further analysis due to nonlinear artifacts that can be introduced due to stimulation application^105-107^. All EEG and physiological data used for further analysis were then bandpass filtered with a 6th order Butterworth filter between 0.25-35 Hz, downsampled to 100 Hz, and min-max normalized.

Behavioral data (CTT circle deviation) were smoothed with a 5 sec moving average window then averaged for the 30 sec pre (D_pre_), and during stimulation (D_during_; **Figure 2j**). The calculated percent change in deviation between the during stimulation (Δ) and pre stimulation period was used as a marker of response to stimulation, whereas this same calculation for the stimulation off periods was used as marker of response to no stimulation. With this configuration a negative delta (-Δ) indicated that participants’ behavioral performance or response increased or got better with the given condition, whereas as a positive delta (+Δ) indicated that participants’ behavioral performance or response decreased or got worse with the given condition.

#### Input and Labels

To demonstrate our approach, we framed our deep learning model development as a data classification problem, where, giving input EEG and physiological data, models were tasked with predicting whether participants would increase or decrease their behavioral performance (response) in the following 30 secs, with a given stimulation condition (No Stimulation, F30, M30), as compared to the prior 30 secs (**Figure 2j**). Here, models were developed for each stimulation condition (No Stimulation, F30, M30) then compared post prediction using a set of decisions rules and model confidences resulting in a *responsiveness* comparison.

Data from Experiment 2 was exclusively used as model inputs in order to unionize stimulation intervention timing as well as stimulation condition and amplitude distribution. Experiment 2 also consisted of participants who repeated their experimental sessions on different days. These repeated experiments were from participant 12 (repeated participant ID: 19), participant 15 (repeated participant ID: 18), participant 21 (repeated participant ID: 25,26), and participant 22 (repeated participant ID: 23, 24). In this case input data were divided into three main groups: No Stimulation, F30, and M30. This division resulted in 68 trials of no stimulation, 20 trials of F30, and 20 trials of M30, for each session (30 sessions total for Experiment 2).

The minimally processed EEG and ECG data were used as model inputs whereas the binarized percent change in behavioral performance (CCT deviation, ±Δ) was used as trial labels. EEG data were further subsampled from 32 to 14 scalp electrodes. These included EEG channels: FPz, Fz, FC1, FC2, Cz, C3, C4, CP2, CP6, P4, POz, O1, Oz, O2 (**Figure 2c-d**). In total 15 channels (14 EEG, 1 ECG) over 30 secs (15 channels X 3000 samples) were used as model inputs for each trial (**Figure 2k**). Labels (CCT deviation, ±Δ) were converted from floating point numbers to integers (0 or 1) and one-hot encoded. Calculated negative delta deviations (-Δ), which indicated an improvement in behavioral performance with a respective condition or an increase in response, was encoded as a 0; whereas positive delta deviations (+Δ), which indicated a decrement in behavioral performance with a respective condition or a decrease in response, was encoded as a 1.

#### Deep Learning Models and Training

We selected the EEGNet model architecture, a convolutional neural network (CNN) model designed to ingest minimally processed, multichannel EEG data^42^. The model architecture utilizes a sequence of 2D convolutional layers, including depthwise and separable convolutional layers with interleaved batch normalization, dropout^108^, and average pooling (**Figure 2l**). For each stimulation condition the EEGNet architecture was modified in order to reduce the model’s capacity and reduce overfitting^41^.

Two models were developed for each of the three stimulation conditions (No Stimulation, F30, M30). For the No Stimulation models, the number of temporal filters were set to 8, pointwise filters to 16, batch size to 4, learning rate to 5E-5, dropout rate of 50%, and normalization rate of 0.1. The No Stimulation models had kernel sizes of 34 and 64 and were trained for 34 epochs. For the F30 and M30 models, the number of temporal filters were set to 2, pointwise filters to 6, batch size to 30 (100 for M30), learning rate to 5E-3, dropout rate of 80%, and normalization rate of 0.1. Both the F30 and M30 models had kernel sizes of 100 and 105 and were trained for 150 epochs. All models utilized the ADAM^109^ optimizer with focal loss^110^ where alpha and gamma parameters were set to 0.2 and 5.0, respectively.

The training set consisted of participants 12, 14, 16, 18, 20, 22, and 26, this gave a total of 952 trials for No Stimulation (**Figure 3a**); and 140 trials for F30 and M30, respectively (**Figure 3b-c**). The test data consisted of participants 11, 19, 21, 23, 24, 25, with 816 trials for No Stimulation (**Figure 3a**); and 120 trials for F30 and M30, respectively (**Figure 3b-c**). Since training data were comparatively small and the distribution of classes was imbalanced (due to the open-loop effect of stimulation); class weighting and data augmentation were utilized. For data augmentation, normally distributed random noise with 0 mean and standard deviations of 0.25, 0.5, 1.0 and 1.5 were added to the training data, the channels with added noise were then flipped in time, while preserving their channel-wise order^59^.

## Data availability

All raw data acquired and used in the text can be accessed directly at: https://doi.org/10.5281/zenodo.3837212. The dataset is made available in multiple formats across repositories and is compatible across most analysis pipelines including MATLAB and Python. The dataset is additional formatted to the Brain Imaging Data Structure (BIDS) specifications and can be accessed directly at: https://doi.org/10.18112/openneuro.ds003670.v1.1.0. Each stimulation trial within the dataset can be visually inspected in the time and frequency domain directly at https://doi.org/10.6084/m9.figshare.14810517.v1 for Power Spectral Densities, https://doi.org/10.6084/m9.figshare.14810442.v1 for time-frequency spectrograms; and https://doi.org/10.6084/m9.figshare.14810478 for scalp voltage topographies. Additional formatted data is available upon request.

## Code availability

The latest version of all accompanying code for this work is available at: https://github.com/ngebodh/GX_DL_Framework. Additional code to parse and extract aspects of the GX dataset can be acquired within this repository: https://github.com/ngebodh/GX_tES_EEG_Physio_Behavior. Additional resources are available upon request.

## Author Contribution

Nigel Gebodh designed the experiment, collected data, ran data analysis, created data visualizations, and wrote the manuscript. Vladimir Miskovic designed the experiment and wrote the manuscript. Sarah Laszlo designed the experiment and wrote the manuscript. Abhishek Datta designed the experiment and revised the manuscript. Marom Bikson designed the experiment and wrote the manuscript. All the authors conceived the study and revised and approved the final manuscript.

## Acknowledgments

Portions of this study were funded by X (formerly Google X), the Moonshot Factory. The funding source had no influence on study conduction or result evaluation. Marom Bikson is supported by grants from Harold Shames and the National Institutes of Health: NIH-NIDA UG3DA048502, NIH-NIGMS T34 GM137858, NIH-NINDS R01 NS112996, and NIH-NINDS R01 NS101362. Nigel Gebodh and Marom Bikson are further supported by NIH-G-RISE T32GM136499.

We would like to thank Dr. Asif Rahman and Lukas Hirsch for their guidance and input throughout this work.

---

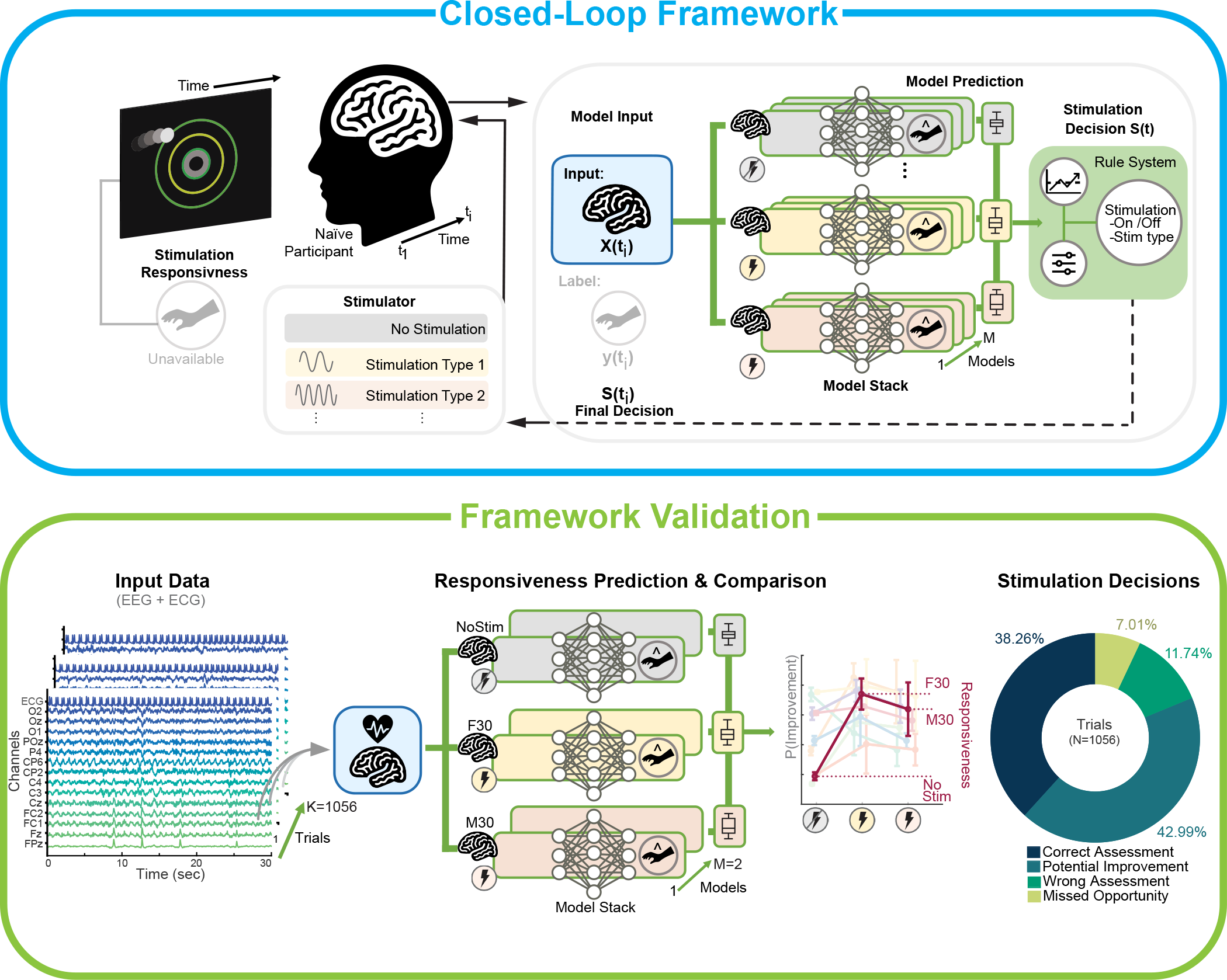

